# An Efficient Computational Approach for Constructing the Allele Frequency Spectrum of Populations with Arbitrary Complex History

**DOI:** 10.1101/456335

**Authors:** Hua Chen

## Abstract

The allele frequency spectrum (AFS), or site frequency spectrum, is commonly used to summarize the genomic polymorphism pattern of a sample, which is informative for inferring population history and detecting natural selection. Recently, Chen and Chen (2013) developed a method for analytically deriving the AFS for populations with temporally varying size through the coalescence time-scaling function. However, their approach is only applicable for population history scenarios in which the analytical form of the time-scaling function is tractable. In this paper, we propose a computational approach to extend the method to populations with arbitrary complex history by numerically approximating the time-scaling function. We demonstrate the performance of the approach by constructing the AFS for two population history scenarios: the logistic growth model and the Gompertz growth model, for which the AFS are unavailable with existing approaches.

## 1. Introduction

The allele frequency spectrum (AFS, a.k.a. the site frequency spectrum) is a series of fundamental statistics for summarizing genomic polymorphism. It is defined as the sampling distribution of allele frequencies of genetic polymorphism in a finite sample (Chen, 2012). In practice, the AFS can be the number or proportion of SNPs constructed by binning them according the counts of derived alleles. For a sample of *n* sequences with *m* identified segregating sites (polymorphic sites), the AFS is written as {(*S_i_*),1 *< i < n*}, with 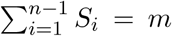, where *S_i_* denotes the number of segregating sites in the sample that has *i* copies of derived alleles among the *n* haplotypes. The AFS has been a main focus in theoretical and methodological studies in the past decades, since it is informative for the inference of ancient demography of populations (Kimura, 1955). The theoretical expectation of AFS under a given population history and parameter setting can be developed using both coalescent theory and diffusion (Fu, 1995; Griffiths and Tavaré, 1998; Sawyer and Hartl, 1992). Methods for ancestral inference based on the AFS are then developed in a Poisson random field framework by assuming each entry of the AFS follows a Poisson distribution with the mean equal to the theoretical expectation of AFS given a population genetic parameter setting (Sawyer and Hartl, 1992; Bustamante *et al.*, 2001; Fu, 1995; Griffiths and Tavaré, 1998; Wooding *et al.*, 2002; Polanski and Kimmel, 2003; Marth *et al.*, 2004; Williamson *et al.*, 2005; Gutenkunst *et al.*, 2009; Lukić *et al.*, 2011; Živković and Stephan, 2011; Chen, 2012; Excoffier *et al.*, 2013; Gao and Keinan, 2015; Bhaskar *et al.*, 2015; Liu and Fu, 2015). These methods gain popularity with the abundance of genomic sequencing data.

Coalescent theory has been applied to developing the AFS in a single population with time-varying population sizes, including the exponential-growth model (Wooding and Rogers, 2002; Polanski and Kimmel, 2003) and the n-epoch model, which models the population size changes using several consecutive periods (epochs) with different constant sizes (Marth *et al.*, 2004). Compared with the AFS developed with diffusion, the coalescent-based AFS has the advantage of being in analytical form, and the estimation is fast and accurate for small samples. In contrast, the diffusion approximation has to rely on numerical methods, such as finite difference approaches, to approximate the solutions (Williamson *et al.*, 2005; Evans *et al.*, 2007). The coalescent-based AFS is thus very useful in the inference of past demographic history and has been extensively applied to data analysis (Marth *et al.*, 2003; Keinan *et al.*, 2007; Gravel *et al.*, 2011; Gazave *et al.*, 2014).

One limitation of the coalescent-based AFS methods is that we can only analytically derive the AFS for some simple population growth models, such as the n-epoch model and the exponential-growth model or their combinations (Polanski and Kimmel, 2003; Marth *et al.*, 2004; Gazave *et al.*, 2014), and generalization to other complex population histories is often impracticable (Chen, 2012, 2013; Polanski and Kimmel, 2003). A second limitation is that for large samples (e.g., haplotype number *n >* 50), it is hard to accurately calculate the expected AFS from the formulae. The expected coalescence times 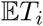, 1 ≤ *i* < *n* are essential for deriving the coalescent-based AFS, which contain coefficients in the alternating sum of the hypergeometric series and are explosively large, causing overflow for large sample sizes (Polanski *et al.*, 2003). When the sample size is large, the AFS and its derived statistics are informative for inferring recent population history. And thus, calculating the AFS for large samples becomes common in population genetic inference from genomic data (Coventry *et al.*, 2010; Gazave *et al.*, 2014; Chen *et al.*, 2015). A high-precision arithmetic library is usually adopted to obtain accurate numerical values when analyzing larger samples, which requires tedious programming and intensive computation (Marth *et al.*, 2004). Some alternative solutions were proposed, specifically for the AFS of a single population, e.g., Polanski and Kimmel (2003) replaced it with hypergeometric summation to avoid estimating the coefficients with large values. Their approach can efficiently solve the numerical issue, but it is difficult to generalize this approach to other scenarios with complex population histories for which the integral function in the hypergeometric summation is difficult to compute. Most studies have adopted coalescent simulations to generate a large samples to approximate the AFS under specific demographic histories and applied them to analyzing genomic polymorphism. However, this approach is computationally very intensive (Hudson, 2002; Coventry *et al.*, 2010; Nelson *et al.*, 2012; Excoffier *et al.*, 2013; Gazave *et al.*, 2014; Tennessen *et al.*, 2012).

To address the numerical issue in large samples, Chen and Chen (2013) used the large-sample asymptotic distributions of coalescence times. Griffiths (1984) proved that the coalescence times and ancestral lineage numbers asymptotically follow a normal distribution in a constant population. Chen and Chen (2013) extended their forms to populations with time-varying sizes by using a time-scaling function scheme (see the “Coalescence times” section below; Griffiths and Tavaré (1994); Donnelly and Tavaré (1995); Nordborg (2001)) and then used the first-order Taylor expansion approximation to achieve the coalescence times (and, further, the AFS). They illustrated the usage of this approach by deriving a simple-form formula for the AFS in populations under exponential growth, which shows high accuracy compared with simulated results. Note that the first-order Taylor expansion approximation and time-scaling function approach of Chen and Chen (2013) works for both large and small size samples. Technically, their approach allows them to derive AFS in any populations with arbitrary complex demography. However, as we illustrate in the “Methods” section, for some complex demography, it is difficult to derive the analytical form of the time-scaling function and/or its inverse function, which are essential in deriving the coalescent-based AFS. In this paper, we propose a computational approach to efficiently approximate the analytical formula of the time-scaling function with a finite sum approximation, and find the set of coalescence times 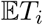, 1 ≤ *i < n*, with the computing time being nearly constant as the sample size increases. It is applicable to any arbitrary complex history for which the time-scaling function is not tractable. This greatly extends the application of AFS-based methods in population genetic inference and other studies, e.g., cancer evolution. We demonstrate the performance of the approach by obtaining the AFS for two population history scenarios that were difficult to derive using the existing approaches: the logistic growth model and the Gompertz model.

In the following sections we first review the coalescent theory framework for obtaining the AFS for a single population. We then summarize the first-order Taylor expansion approximation method for populations with time-varying size proposed by Chen and Chen (2013). We illustrate the idea of the computational approach for constructing the AFS for arbitrary demography, and we further derive the AFS for populations with two demographic histories to demonstrate its performance.

### Modeling framework

For a sample of *n* lineages (haplotypes), the coalescence time *T_k_* is defined as the time when *k* + 1 lineages merge into *k* lineages, and time is measured backward (from the present to the past). The intercoalescence time *W_k_* = *T_k_*_−1_ − *T_k_* is the time during which there are *k* lineages. Following Fu (1995), we say that any of the *k* branches spanning the intercoalescence time *W_k_* has the branch of size *k*. We assume an infinitely-many-sites model for mutations, and further mutations occur on branches along the gene genealogy following a Poisson process. The number of mutations occurring at any branch of size *k* then follows a Poisson distribution with the mean of 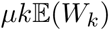, where *µ* is the point mutation rate. During the bifurcation process in which *k* lineages increase to *n* lineages at present, any of these mutations increases the allele account from a single copy to *j* among the *n* lineages with the probability (Feller, 2008; Griffiths and Tavaré, 1998):

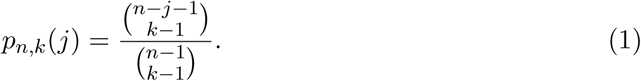

Summing over mutations that occur on branches with different sizes, we can obtain the entries for the AFS:

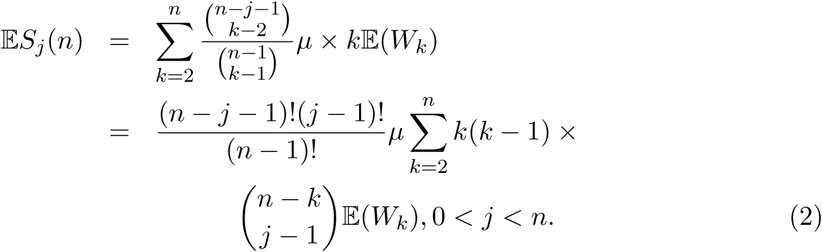

Note that 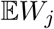 is fundamental in the above framework for constructing the AFS. If we can obtain analytical forms for 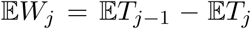 for a population with complex demography, we can obtain the AFS through Equation 2.

### Coalescence times

In a constant-size population, the distribution of coalescence times follows that of the standard Kingman’s n-coalescent, which are exponential variables with the mean

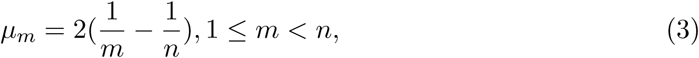

where *µ_m_* is the coalescence time in units of haploid population size *N*. In addition, the intercoalescence times are mutually independent.

For a population with time-varying size, we denote its population history as *N*(*t*), *t* ∈ [0, ∞). It is not trivial to derive the distribution or the expectation of coalescence times for a population with time-varying sizes. The joint distribution of coalescence times (*T_m_*, …, *T_n_*_−1_) for populations with time-varying size is (Griffiths and Tavaré, 1998)

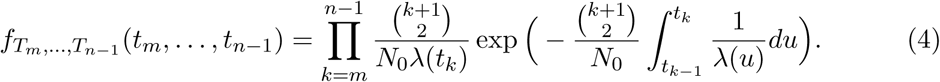

Polanski *et al.* (2003) derived the marginal probability density function of coalescence times 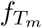 by expanding an integral transform of the marginal pdf into partial fractions. Another way to derive 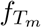 is based on the definition of a pure-death process, in the form of a function of the ancestral lineage number, *P*(*A_n_*(*t*) = *m*) (Griffiths, 2006; Chen, 2012). With the marginal distribution of coalescence times derived, Polanski and Kimmel (2003) obtained the AFS for a population under exponential growth, which is in complex form, and requires calculating the hypergeometric series and exponential integral.

Chen and Chen (2013) used the time rescaling approach in the variable-population-size coalescent model (Griffiths and Tavaré, 1994; Nordborg, 2001; Donnelly and Tavaré, 1995). The coalescence time is rescaled at the rate 1/*N*(*t*), denoted as *τ_m_*:

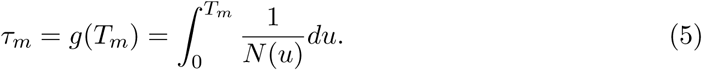

*τ_m_* follows the coalescence time distribution in the standard Kingman’s n-coalescent (Kingman, 1982). Chen and Chen (2013) then used a Taylor expansion of *T_m_* = *g*^−1^(*τ_m_*) around *µ_m_* to achieve the approximation:

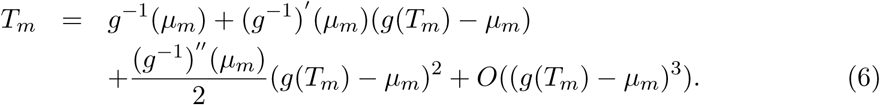

Thus

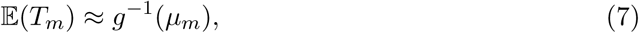

and

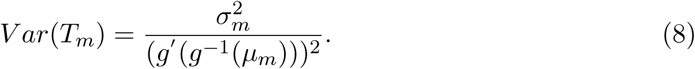

In general, for any population history *N*(*t*), 0 ≤ *t* < ∞, we can always obtain the time-scaling function *g*(*t*) as in Equation 5, and further obtain 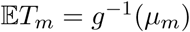 as above. Chen and Chen (2013) demonstrate the application of this approach using an exponentially growing population as an example. 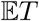 for the exponential growth model is in a very simple analytical form:

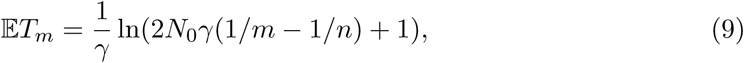

and the obtained AFS is highly accurate (see Figure 6 of Chen and Chen (2013)).

Since it is not trivial to derive the coalescence times for populations with time-varying size in existing studies, and simulations are usually required as a replacement for most studies, Chen and Chen (2013)’s approach provides simple and efficient solution for obtaining 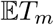 (Coventry *et al.*, 2010; Nelson *et al.*, 2012; Tennessen *et al.*, 2012; Excoffier *et al.*, 2013; Gazave *et al.*, 2014). However, for some complex demographies, the analytical form of the time-scaling function *g*(*t*) and its inverse function, which are essential for deriving 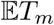, are not tractable. This prohibits the general usage of their approach for arbitrary population history.

### Coalescence times under complex demographic history

In this section, we illustrate how to extend Chen and Chen (2013)’s method to be applicable to arbitrary population history using a computational approach. As we can see from the above section, *g*(*t*) and *g*^−1^(*t*) are the two essential components for deriving coalescence times for a given population history *N*(*t*) (see Equation 7). Note that to obtain 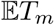, we do not need the analytical form for calculating an arbitrary point *t.* In contrast, we only need to find a finite number of *T_m_* values that correspond to *µ_m_*, 1 ≤ *m* < *n* and satisfy

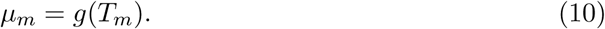

We thus propose the following two numerical schemes for calculating 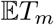, applicable to different situations. The first approach is generally applicable to all cases, including those for which we cannot obtain *g*(*t*); the second approach is specifically for the case in which we have an analytical form of *g*(*t*) but *g*^−1^(*t*) is not tractable.

#### Approach 1 (finite sum approximation)

For a sample of size *n* under the population history *N*(*t*), *t* ∈ [0, ∞), we can simply approximate the integral of the time scaling function equation using the discrete finite summation:

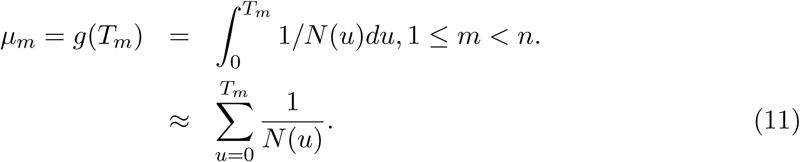

Then, for each *µ_m_*, the corresponding expected coalescence times 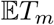 can be obtained during the following sequential summation procedures:

Step 1 We have a series of expected coalescence times under the standard n-Kingman’s coalescent 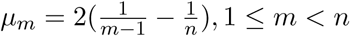. Initialize the procedure from generation 0 (the current generation) with 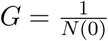.

Step 2 Keep increasing the discrete generation time *t*, and calculate 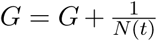 until the value *t* satisfies 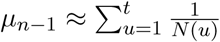. Set *T*_*n*−1_ = *t*.

Step 3 Repeat Step 2, and keep increasing *t* to obtain the rest of the values for 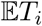, *n* − 2 ≤ *i* ≤ 1.

Step 4 terminate the process when 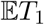 is obtained.

After we have {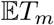, 1 ≤ *m* < *n*}, the AFS can be constructed through Equation 2. The detailed pseudocode for implementing the algorithm is listed in Table 1.

**Table 1:**
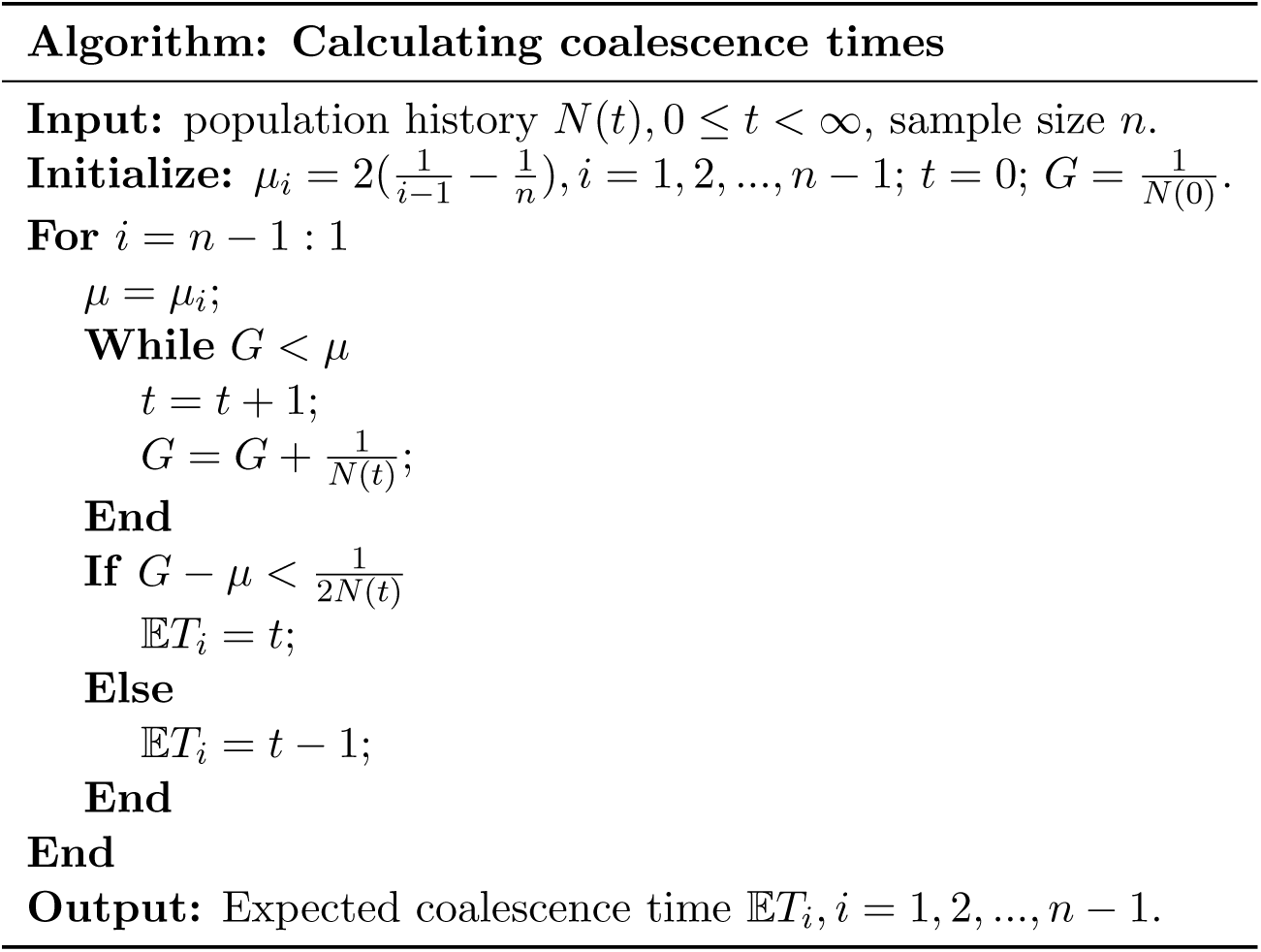
Procedures for calculating coalescence times using the finite-sum approximation (Approach 1).

#### Approach 2

For some population histories, the analytical form of the time scaling function *g*(*t*) can be achieved, but the inverse function *g*^−1^(*t*) is not tractable. An alternative approach can be applied to obtain 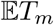 for such cases through the following procedures. For each *T_m_*, 1 ≤ *m* < *n*, we have the non-linear equation,

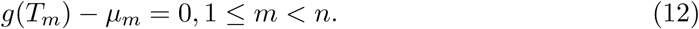

The above non-linear equations can be solved using numerically algorithms to obtain *T_m_*. such as Newton-Raphson (Press *et al.*, 1992). In this paper, we adopt two approaches implemented in MATLAB. The first one is the fzero function, which implements Dekker’s algorithm as a combination of bisection, secant, and inverse quadratic interpolation methods (Brent, 2013). The second is the fminsearch function, which uses the simplex search method of Lagarias *et al.* (1998)

This approach usually takes more time than Approach 1, as for each coalescence time *T_m_*, we need to solve the corresponding equation iteratively. Furthermore, the number of equations and the computational complexity increase with the sample size, and thus Approach 2 is more suitable for small samples.

## Results

Various population growth models have been proposed to approximate the ancient population history of humans and other species. For example, Gazave *et al.* (2014) proposed a five-scenario model for the European population, including two stages of population bottlenecks and a very recent exponential growth. The simple exponential population growth model may be the most commonly used model. It assumes a constant growth rate, which is valid when space and resources are unlimited. The exponential growth model is a good approximation for the early stage of humans, bacteria, and most populations. In cancer evolution studies, models with more parameters were developed to describe tumor growth (Benzekry *et al.*, 2014). These models are complicated by modifying growth rates with carrying capacity or other factors, e.g., the logistic growth model and Gompertz model.

In this section, we use exponential, logistic and Gompertz growth models to illustrate the usage of our proposed approach. For the exponential growth model, *N*(*t*) = *N*_0_*e*^−*rt*^, it is straightforward to analytically derive the expected coalescence times 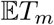 (Equation 9). We then compared the running time of the three approaches (including the analytical approach, the finite-sum approximation, and Approach 2) for the model with the two parameters *N*_0_ = 100, 000 and *r* = 0.003. For Approach 2, we adopted two numerical methods: the bisection + interpolation method implemented in the MATLAB function fzero and the downhill simplex method implemented in the MATLAB function fminsearch. The time was averaged over 1000 repeats running in MATLAB and is recorded in Table 2. The detailed results for the logistic growth and Gompertz model are elaborated below.

**Table 2:**
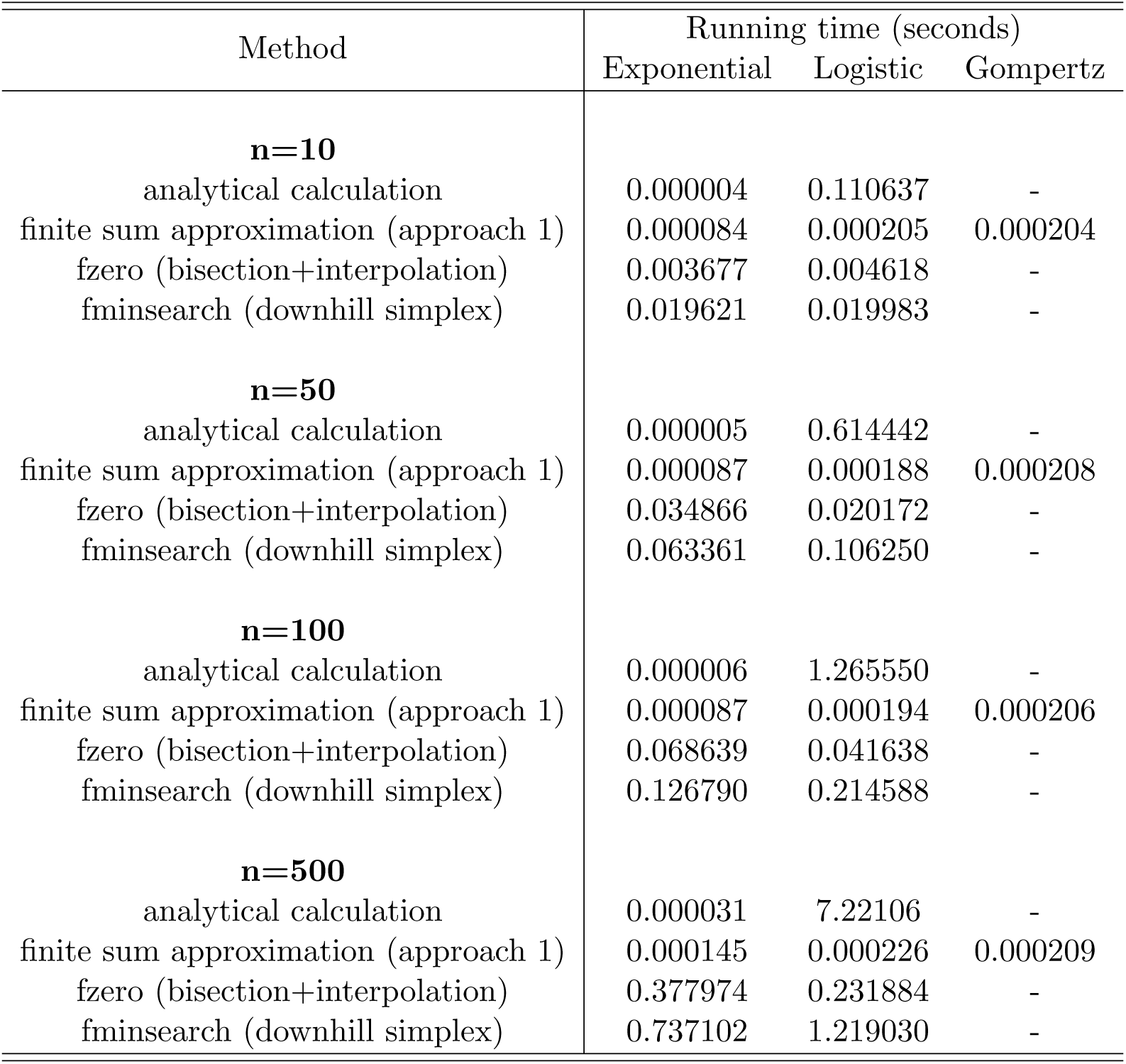
Comparison of running times between different methods for three population growth models and four different sample sizes (10, 50, 100 and 500). For the Gompertz model, only the results of the finite-sum approximation are available.

### Logistic growth

Compared with the exponential growth model, the logistic growth model regulates the growth rate with a factor 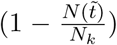, in which *N_k_* is the carrying capacity. It thus has a sigmoid shape and reaches an equilibrium size of *N_k_* instead of unlimited growth (see Figure 1(A)). A logistic growth model is consistent with the population dynamics of many organisms and is widely used in ecological research. Let *γ* be the maximum population growth rate (aka, intrinsic growth rate); for a population under logistic growth,

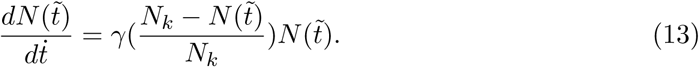

**Figure 1:**
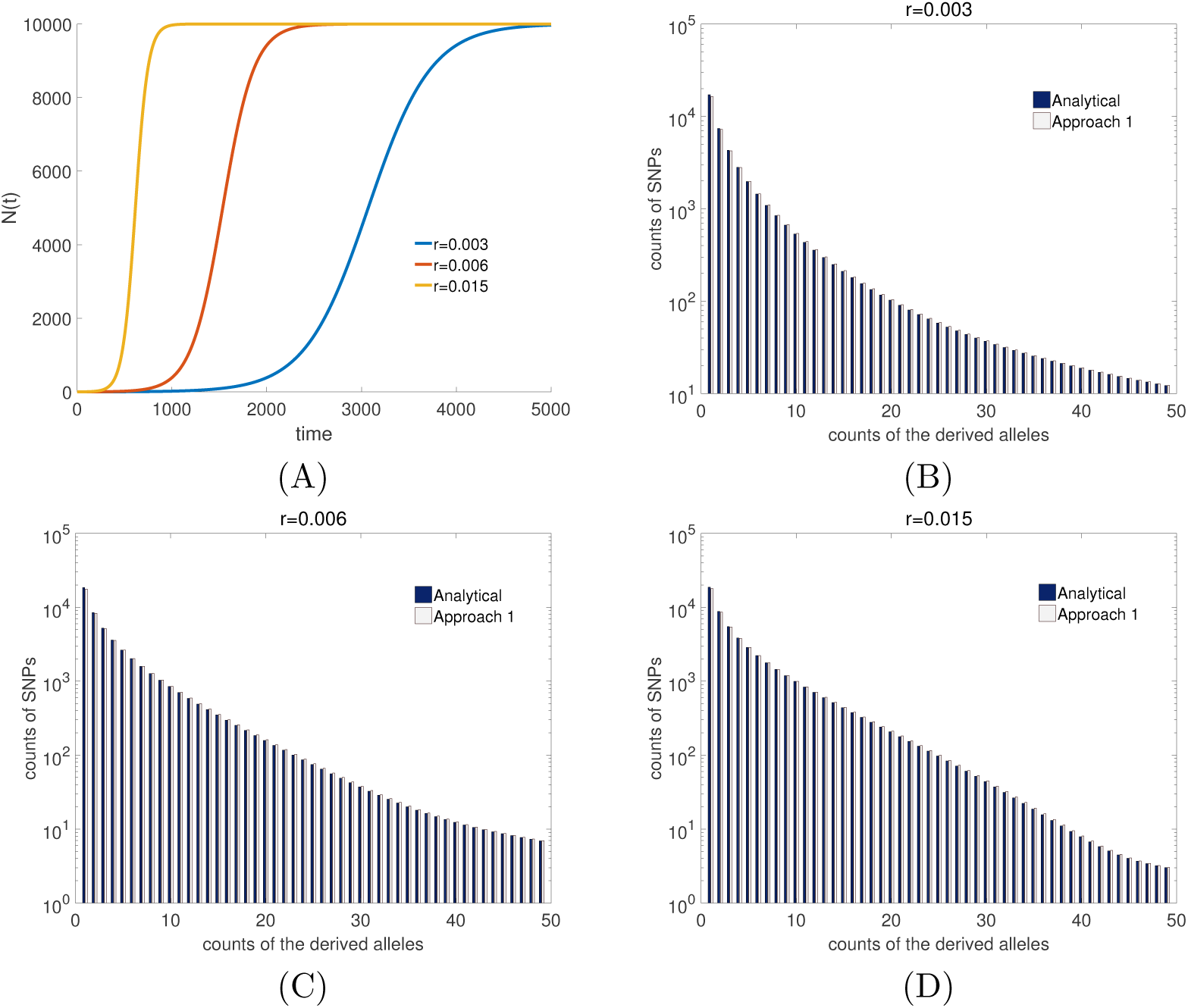
The population growth rate (A) and population size (B) as a function of time, and the allele frequency spectrum (C) of the logistic growth model for three growth rates: *γ* = 0.003, 0.006 and 0.015. The other parameters are: *N_k_* = 10, 000 and *T* = 5000.

Note that in the above equation, time is measured forward (from the past to the present), and we denote it with 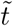 to distinguish it from the backward time in other sections. The population size 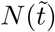 follows a logistic curve,

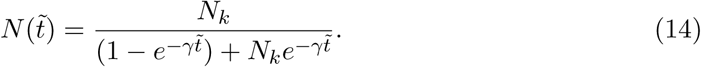

After changing the variable of forward time 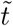 to backward time *t*, we have

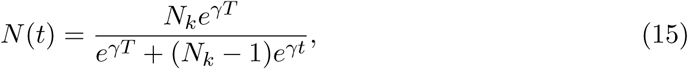

and the model includes three free parameters: *N_k_*, *γ* and *T*.

Given the population history function *N*(*t*), we can derive the time-scaling function for the logistic growth model,

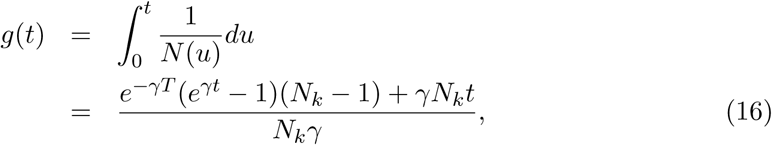

and we further obtain its inverse function,

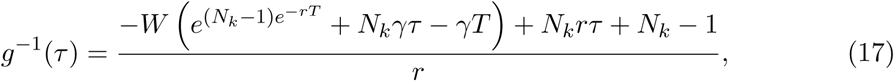

where *W*(·) is the Lambert W function, which is calculated numerically.

According to Chen and Chen (2013), the expected coalescence time 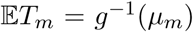 can be obtained from Equation 17. We can also calculate it through Approaches 1 and 2 of the previous section. In Figure 1, we present the AFS generated from Equation 17 (“Analytical calculation”) and Approach 1 (“Approach 1”) for *N_k_* = 10, 000, *T* = 5000 and three different growth rates *γ* = 0.003, 0.006 and 0.015. We also obtained the AFS using Approach 2 and compared the running time for a specific parameter setting (*N_k_* = 10, 000, *T* = 5000, and *γ* = 0.005) for three approaches (Table 2).

### Gompertz growth

The Gompertz model is another widely used model to approximate population dynamics. It was originally proposed to explain human mortality (Gompertz, 1825) and is also used to describe the population growth of other species, including bacteria, animals, and plants (Tjørve and Tjørve, 2017). The Gompertz model was found to fit well the growth of breast cancer and 19 other tumor cell populations (Laird, 1964; Norton *et al.*, 1976; Norton, 1988). One of its forms is

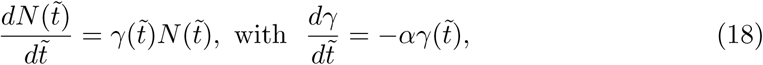

where *γ*_0_ is the initial growth rate; *N*_0_ is the initial population size when it started to grow; and *α* can be viewed as the exponentially decaying rate of the growth rate. The population history 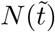 is determined by four parameters: *N*_0_, *γ*_0_, *α* and *T*, the duration since the population growth began. The solution of the above differential equation is

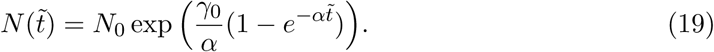

It is unfeasible to derive the time-scaling function *g*(*t*) and its inverse function *g*^−1^(*t*) for the Gompertz model. Therefore, we have no analytical calculation or numerical solution (Approach 2) of the coalescence times for the Gompertz model. In Figure 2 (A) and (B), we show the growth rates and population size trajectories as a function of time for four parameter settings: *γ*_0_ = 0.01, *α* = 0.0005; *γ*_0_ = 0.01, *α* = 0.001; *γ* = 0.05, *α* = 0.004; and *γ* = 0.05, *α* = 0.008. The corresponding AFS for *n* = 50 haplotypes is presented in Figure 2 (C)-(F). The running times of Approach 1 for a specific parameter setting (*T* = 5000, *r* = 0.01, *α* = 0.001 and *N*_0_ = 1) and with different sample sizes (10, 50, 100, and 500) are presented in Table 2.

**Figure 2:**
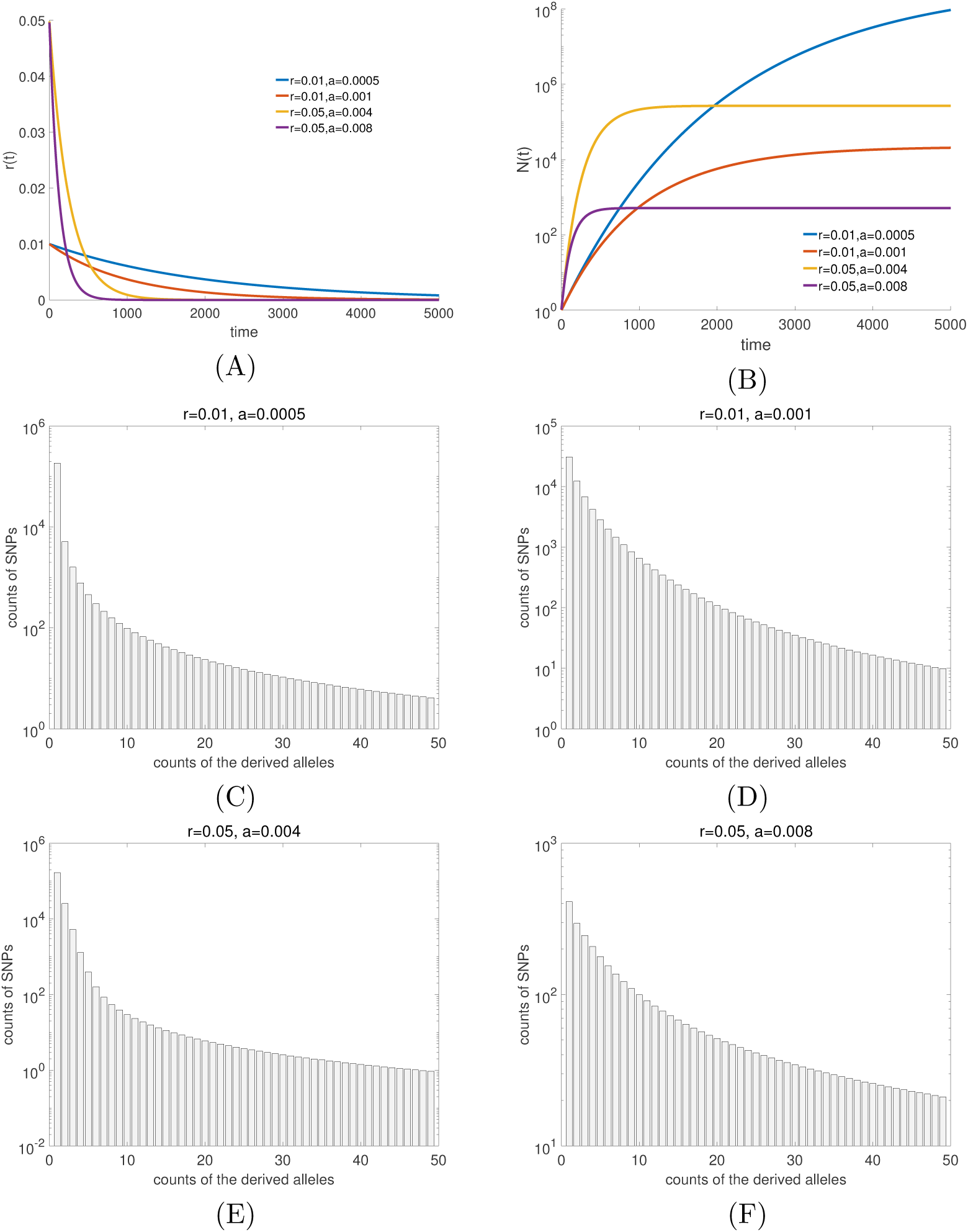
The population growth rate (A) and population size (B) as a function of time, and the allele frequency spectrum (C) of the Gompertz model for four parameter settings: *γ* = 0.01, *α* = 0.005; *γ* = 0.01, *α* = 0.001; *γ* = 0.05, *α* = 0.004 and *γ* = 0.05, *α* = 0.008. The other parameters are: *N*_0_ = 1, *T* = 5000.

### Computing time

We compared the computing times for Approach 1 (finite-sum approximation), Approach 2 and the analytical approach. For Approach 2, we used several methods for solving the non-linear equations, including the bisection+interpolation (implemented in the MAT-LAB function fzero) and the downhill simplex (implemented in fminsearch) methods. All the comparisons are in MATLAB. We investigated three population growth models: the exponential growth, logistic growth and Gompertz growth model. The running times for constructing the coalescence times *T_m_*, 1 ≤ *m < n*, for four sample sizes *n* = 10, 50,100 and 500 were recorded. The running time was averaged over 1000 repeats, as listed in Table 1 (in seconds).

A trend in Table 2 worth noting is that the finite-sum approximation is very fast. The running time is close to that of the analytical calculation, nearly of the same magnitude, and much faster than that of numerical approaches (Approach 2). The only outlier is the logistic model,for which the finite-sum approximation is much faster than the analytical approach. This is because the analytical form of the *g*(*t*) function for the logistic model consists of the Lambert W function, which is calculated numerically and is time-consuming.

Second, note that the running time of the finite-sum approximation approach is nearly constant with increasing sample size *n*. As we mentioned above, the computational complexity is *O*(1), and thus, it is insensitive to the sample size. This guarantees the computational efficiency of the approach when the sample size becomes large, enabling its application to large-sample data analysis.

The numerical approach for solving the *g*(*t*) function (Approach 2) also works efficiently but is more time-consuming than the finite-sum approximation approach for all three population growth models. Furthermore, the running time increases with the sample size *n*, as the number of nonlinear equations to solve increases linearly with *n*.

## Conclusion

The allele frequency spectrum is informative for population genetic inference. Various AFS-based methods have been developed for inferring population history and detecting natural selection in the past years. They have gained popularity with the abundance of genomic sequencing data (e.g., Bustamante *et al.* (2001); Griffiths and Tavaré (1998); Polanski and Kimmel (2003); Marth *et al.* (2004); Nielsen *et al.* (2005); Williamson *et al.* (2005); Gutenkunst *et al.* (2009), etc.). Compared with the diffusion-based AFS methods which require approximation of the solutions with numerical approaches, modeling the AFS using coalescent theory is computationally efficient. Most population genetic inference methods using the coalescent likelihood require computationally intensive algorithms for parameter estimation, such as importance sampling or Markov chain Monte Carlo, while the coalescent-based AFS methods only depend on the expected coalescence times, which guarantee the analytical form (Fu, 1995; Griffiths and Tavaré, 1998; Chen, 2012).

The coalescent-based AFS has shortcomings. First, for large samples it is impossible to obtain accurate calculations due to numerical overflow of large coefficients in the hypergeometric series. Second, it is difficult to derive the coalescent-based AFS for complex population histories, which limits its application to simple growth models, such as the exponential growth and n-epoch models. Chen and Chen (2013) showed that for complex demography, we can obtain the expected coalescence times through a linear Taylor expansion approximation, which involves the time-scaling function g(t) and its inverse function *g*^−1^(*t*). The analytical equations of coalescence times derived through this approach are in a simple form and can successfully overcome the numerical issue for large samples. Furthermore, the time-scaling scheme is technically applicable to arbitrary complex population histories. However, in practice, the analytical forms of the population-scaling function *g*(*t*) and its inverse function are not achievable for many cases, limiting the applications. For example, in the study of cancer cell growth, various population growth models in complex form were proposed to describe the dynamics of cancer cells, for which the analytical form of AFS is difficult to derive. In this paper, we propose a computational approach, the finite-sum approximation, efficiently solving the problem of Chen and Chen (2013) when the analytical form of the time-scaling function *g*(*t*) and its inverse function *g*^−1^(*t*) are not derivable.

We apply the computational approach to three widely used models, including the exponential, logistic and Gompertz growth models to demonstrate its performance. As shown in the Results section, the finite-sum approximation approach is computationally very efficient, and the running time is nearly on the magnitude of that of the analytical approach. Furthermore, the computational time does not increase linearly with the sample size, ensuring its efficiency for AFS of large sample sizes. This is especially attractive for the flexibility to tackle a complex population history that is intractable by using the analytical approach, for example, the Gompertz growth model shown in Table 2. The computational approach presented in this paper is applicable to arbitrary complex population history and significantly enables the application of the coalescent-based AFS approaches to population genetic inference in the genomic sequencing era.

## Acknowledgements

I am are grateful to Kun Chen for helpful discussions and to Shilei Zhao for helping with the numerical methods in MATLAB. This project was supported by the National Natural Science Foundation of China (Grant No. 31571370, 91631106 and 91731302), the Strategic Priority Research Program of the Chinese Academy of Sciences (Grant No. XDB13020400) and the “One Hundred Talents Program” of the Chinese Academy of Sciences.

